# Infection of highly insecticide-resistant malaria vector *Anopheles coluzzii* with an environmentally friendly entomopathogenic bacteria *Chromobacterium violaceum* reduces its survival, blood feeding propensity and fecundity

**DOI:** 10.1101/366484

**Authors:** Edounou Jacques Gnambani, Etienne Bilgo, Adama Sanou, Roch K. Dabiré, Abdoulaye Diabaté

**Affiliations:** Institut de Recherche en Sciences de la Santé (IRSS) / Centre Muraz, Bobo Dioulasso, Burkina Faso; Université Ouaga 1, Prof Joseph Ki Zerbo, Ouagadougou, Burkina Faso; Université Nazi Boni / Centre Muraz, Bobo-Dioulasso, Burkina Faso

## Abstract

This is now a concern that malaria eradication will not be achieved without the introduction of novel control tools. Microbiological control might be able to make a greater contribution to vector control in the future. Here, we studied the impact of *Chromobacterium violaceum* infections isolated from wild caught *Anopheles gambiae* s.l. mosquitoes in Burkina Faso on mosquito survival, blood feeding and fecundity propensy. *C. violaceum* kills pyrethroid resistant mosquitoes *An. coluzzii* (LT80 ~ at 10^8^ bacteria cell/ml of sugar meal). Interestingly, this bacterium had other negative effects on mosquito lifespan by significantly reducing (~59%, P<0.001) the mosquito feeding willingness from day 4-post infection to 9-day post infection. Moreover, *C.violaceum* considerably jeopardized the mosquito egg laying and hatching of mosquitoes by ~77.93% and ~22 % respectively. Mosquitoes infected with *C. violaceum* also showed significantly higher retention rates of immature eggs and follicles. These data showed important entomopathogenic properties of Burkina Faso *C. violaceum strains*. However, additional studies as the sequencing of *C. violaceum* genome and the potential toxins secreted will certainly provide useful information render it a potential candidate for the biological control strategies of malaria.

## Background

In the last three years, many countries have reported significant increases in malaria cases, according to the WHO new malaria report [1]. Progress toward the global elimination of malaria is then stalling. In 2016, the number of malaria cases worldwide was ~216 millions [1]. That is down from 237 millions in 2010, but about the same as it was been the last few years [1–3]. With 194 million cases in 2016, Africa accounted for almost 90% of the total. The estimated number of deaths was ~ 445,000 from malaria in 2016. More than 90% of those deaths were in Africa, and 80% of the victims in Africa are children younger than 5 and pregnant women [1]. The reasons for the slowdown differed across specific regions and countries. But contributing factors included insufficient funding, a lack of interventions to prevent spread of the disease, risks posed by conflict in malaria endemic regions, irregular climate patterns [4], and the rapid emergence of both parasite and mosquito vector resistance to drugs and insecticides [1]. Emergence and spread of resistance to pyrethroids, organophosphates and carbamates is a particular threat, as most malaria control programs rely heavily on these broad-spectrum chemical insecticides to reduce vector populations [5, 6].

These outgrowing problem drove the WHO to call for researching and developing alternative approaches in controlling vector-borne diseases, thus decreasing the usage of insecticides [7]. Integrated vector management (IVM) efforts are now oriented towards controlling Anopheles either at the larval stages and/or at the adult stages using means of microbial control namely fungi and bacteria, where various concerns at the ecological, environmental, social, and economical levels are highly considered [8, 9]. Many employed approaches and future currently set plans are now focusing on the use of genetically engineered or nor microorganisms to either block the development of the malaria parasite within the Anopheles vector [10–14], or target the vector itself [10, 15]. Despite intensive efforts to develop entomopathogenic bacteria as biocontrol agents against malaria vectors, the strains under investigation have not met expectations due to some functional and practical limitations [8, 9]. Bacteria as *Bacillus thuringiensis israelensis* (Bti) and *Bacillus sphaericus* (Bs) infections show no residual persistence post application [8].Only few studies address the effect of different bacterial agents on malaria vectors [9]. Most of these studies are only experimentally approached without any further practical applications [8]. Regarding the promising entomopathogenic bacteria Wolbachia, only a strain *(wAnga)* is know to be associated with *Anopheles gambiae* [16]. Most of promising *Wolbachia* strains were not found to naturally infect and colonized Anopheles mosquitoes [8, 9].

Interestingly, in a very recent study, Ramirez and collaborators in 2014, showed that a *Chromobacterium sp*. isolate, *Csp_P*, previously isolated from the midgut of field-collected *Aedes. aegypti* mosquitoes, has unique properties which can kill larvae and adults of multiple mosquito vector species, and exerts in vitro anti-Plasmodium and anti-dengue virus activity suggesting that it could be a highly potent candidate for developing weapons against current and future mosquito-borne diseases [8]. To our knowledge Ramirez and collaborators study and other studies have not yet evaluated other properties of Chromobacterium infection that might also contribute to reducing malaria transmission potential such as the blood feeding willingness and the fecundity. In this study, we examined the impact of infection of malaria vector *(An. coluzzii)* with an indigenous burkinabè environmentally friendly strain of *Chromobacterium violaceum* isolated from both wild caught adults and larvae of *Anopheles gambiae*. Through logistically simple method of infection, cotton balls soaked with sugar meal containing bacteria we specifically, assessed the pathogenicity of this local strain of *C. violaceum* against adult mosquitoes and its impact of mosquito blood feeding and fecundity propensy.

## Materials and methods

### Bacteria strain

*Chromobacterium violaceum* strain used for bioassays is isolated from both field collected larvae and cuticles of adult mosquito *(Anopheles gambiae s.l.)* from Bana (11° 9’ 41"N, 4° 10’ 30" W), Soumousso (11°04’ N, 4°03’ W) in western Burkina Faso. Homogenates from dead mosquitoes were firstly plated out onto Chocolate + polyvitek agar and Bromocresol purple agar. Then, 24 hours and 48 hours, bacterial species were isolates and species of *C. violaceum* were identified using the VITEK2 system (**Supplementary file 1)** in laboratory at Centre Muraz.

*C. violaceum* is a rarely human pathogenic Gram-negative facultative anaerobic, non sporing-coccobacillus. It is part of the normal flora of water and soil of tropical and sub-tropical regions of the world. It produces a natural antibiotic called violacein [17]. Some strains of Chromobacterium have already been developed for agricultural pest control [18].

### Mosquito colonies and PCR determination of Kdr levels

For bioassays, we used F1 progeny of *An. Coluzzii* reared from larval collections at Kou Valley (11°23’ N, 4°24’ W). Mosquitoes from these areas are highly resistant to multiple insecticides [19]. Only non-blood-fed females, 2 to 5 days old, were used in bioassays. All tests were carried out at 25 ± 2°C and 80±10% relative humidity.

The level of kdr resistance within a subsample of mosquitoes was performed using the PCR protocol and primer sequences previously described [20]. We only analyzed mutation L1014F because it is the commonest in West Africa, whereas the L1014S mutation is confined to East Africa [19]. The primers AgD1 (5’-ATA GAT TCC CCG ACC ATG-3’) and AgD3 (5’-AAT TTG CAT TAC TTA CGA CA-3’) amplified the resistant allele yielding 195 bp fragments. The susceptible allele was assayed using primers AgD2 (5’-AGA CAA GGA TGA TGA ACC-3’) and AgD4 (5’-CTG TAG TGA TAG GAA ATT TA-3’), which amplified a 137 bp fragment. The primer set AgD1 and AgD2 amplified a ubiquitous 293 bp fragment as a positive control. During amplification, denaturation was set at 94 °C for 3 min followed by 35 cycles of denaturation, annealing and elongation (94°C for 30 s, 55°C for 30 s, 72°C for 10 s, respectively). The final elongation was set at 72 °C for 5 min.

### Bacterial Infection formulation

Mosquitoes used for bioassays were not antibiotic treated. They were maintained on 6% glucose for 2–5 days post emergence. Mosquitoes were then starved overnight and fed for 24 h on cotton balls moistened with a 6 % glucose solution containing *C. violaceum* at desired concentrations (bacterial cells /ml) regarding the purposes of the bioassays (Figure 1). The numbers of bacterial cells were determined by counting through improved Neubauer hemocytometer.

**Figure 1:**
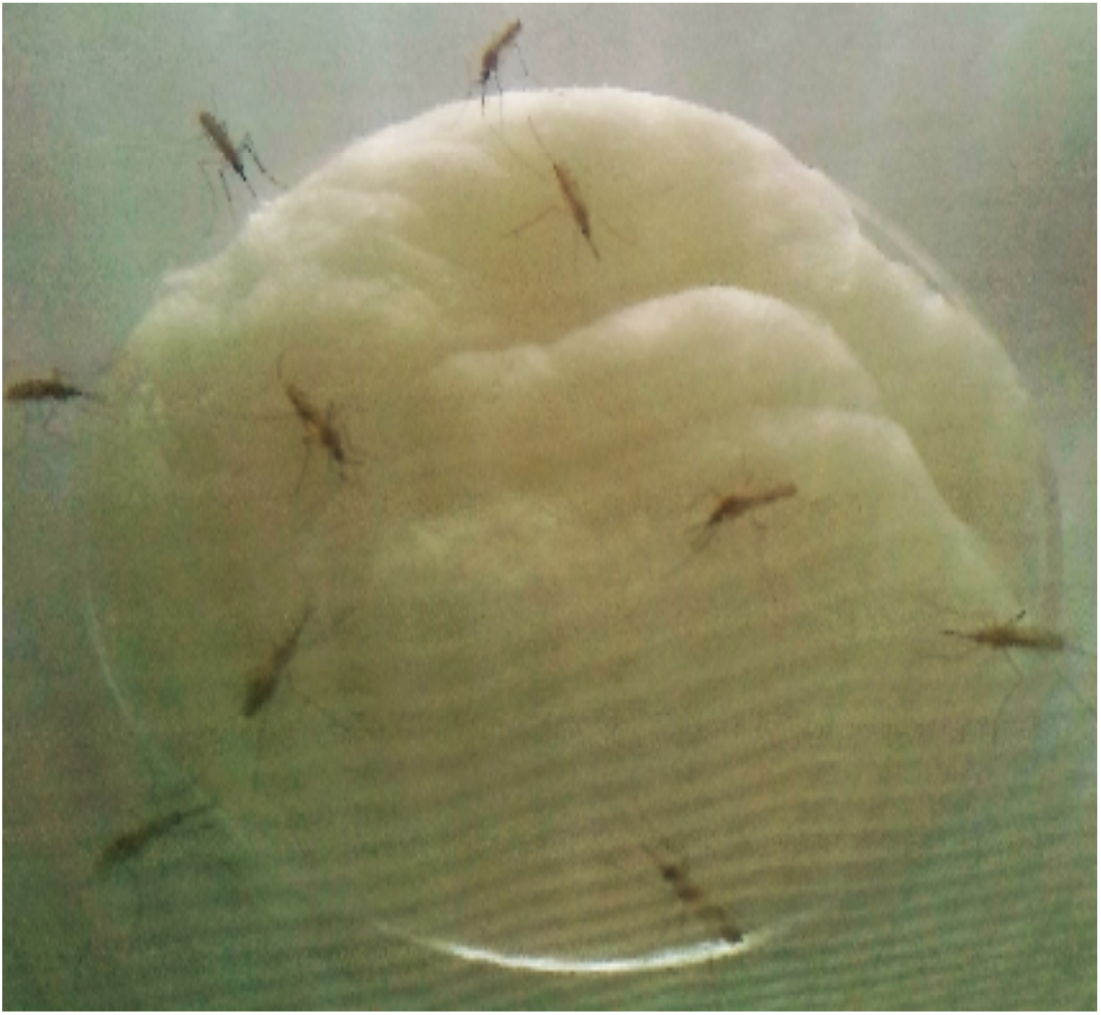
*An. Coluzzii* mosquitoes feeding upon a cotton ball soaked with 6% glucose containing *Chromobacterium violaceum*.

### Bioassays

#### Bioassay1: Evaluation of *C. violaceum* entomopathogenic activity upon mosquito ingestion

To assess the virulence of *C. violaceum* against adults *An. Coluzzii*, 400 newly emerged females were exposed to feed upon cotton balls moistened with a 6 % glucose solution containing 10^5^,10^6^,10^7^ and 10^8^ bacterial cells / ml in a cage (15cm×15cm×15cm) for 24 hours. Control batches of mosquitoes were exposed to blank cotton balls soaked with 6% glucose (without any bacterial cells)

After 24 hours of exposure to treated or untreated cotton balls, mosquitoes were transferred to other new cages (15 cm×15cm×15cm) and fed with sterile 6% glucose. We recorded mortality daily over two weeks. Cadavers were immediately removed from their cages and each was washed once for 20 s with 1% sodium hypochlorite and twice with sterile distilled water for 40 s. Washed cadavers were individually crushed in 200 μl of sterilized phosphate saline buffer (PBS), homogenized. Hundred microliter (100 μl) of each homogenate was then plated onto 2 different media (Chocolate + polyvitek agar and Bromocresol purple agar). Infection was then confirmed 48 hours after incubation using VITEK2 system.

#### Bioassay 2: Blood feeding choice tunnel to assess the impact of *C.violaceum* infection on mosquito host seeking blood-feeding propensy

Fifty non-blood-fed infected or uninfected female mosquitoes (*An. coluzzii*) for each of the four replicates of the tunnel test were released into the tunnel to evaluate the impact of *C. violaceum* infection on mosquito host seeking blood feeding propensy. The bacterial dose (10^6^ bacterial cells / ml) was used for the tunnel bioassay. This dose is suboptimal for killing *An. Coluzzii*, with resulting LT50 of 9 days post infection. Three to nine days post infection mosquitoes were used for the bioassays. The control mosquitoes were infected with blank distilled water solution without any bacteria. We followed the protocol described by Bilgo *et al*. in 2017 [21] with some modifications. The tunnel is basically a 60-cm long; glass choice tunnel (25 cm × 25 cm area) was used for blood feeding assays (Figure 2).

**Figure 2:**
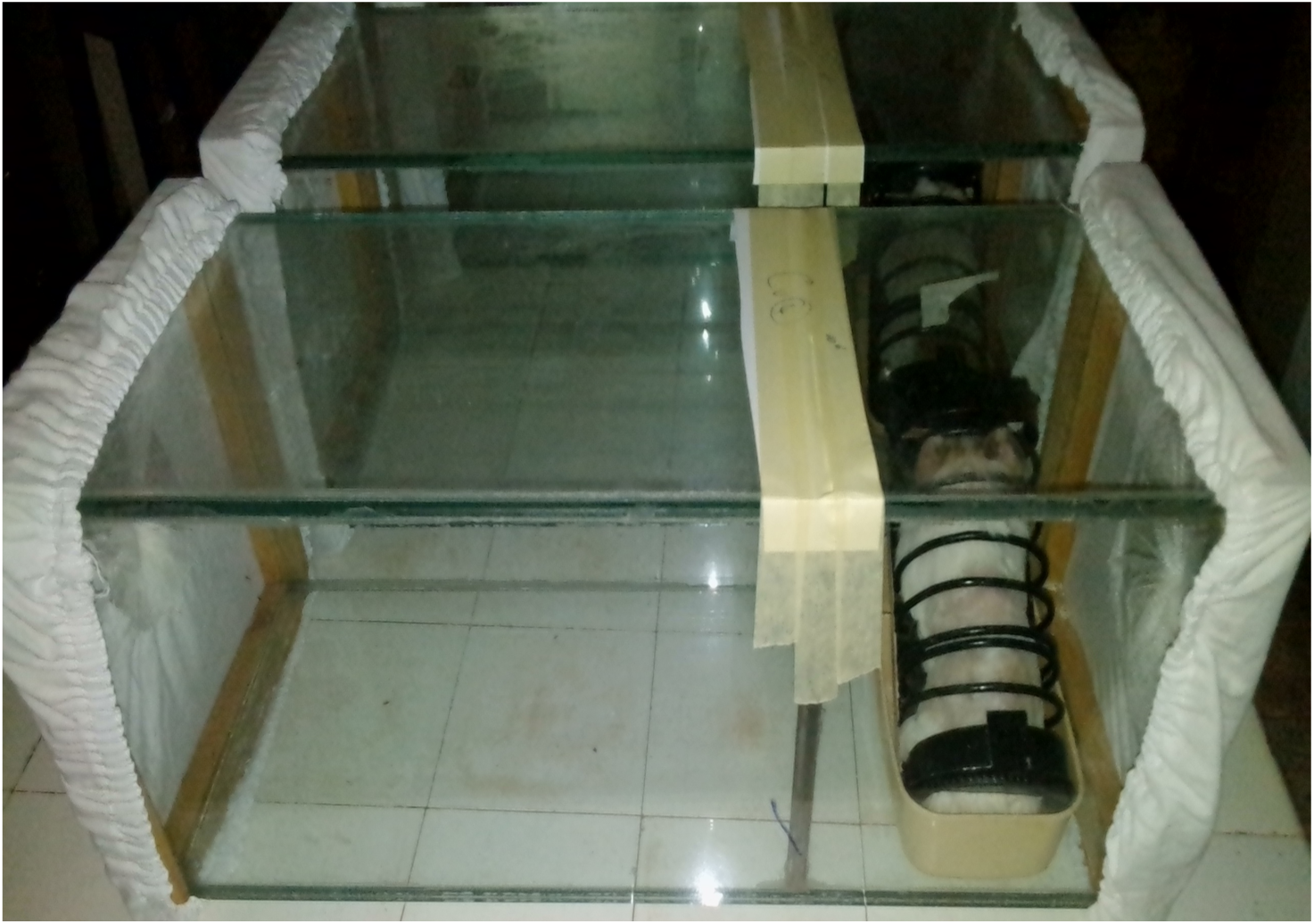
Design for experiments testing host-seeking behavior using guinea pigs and a tunnel choice chamber with nine small holes cut into a barrier between compartments

A 25-cm square of polyester netting was fitted at one end of the tunnel as a compartment. A netted barrier was placed one-third along the length of the glass tunnel separating the tubes into short and long sections. The barrier was 400 cm^2^ (20 × 20 cm), with nine 1 cm diameter holes for passage; one hole was located at the center of the square and the other eight were equidistantly located 5 cm from the border. This choice chamber is designed as a miniaturized proxy for a traditional West African home. The largest section of such a home is the veranda that serves as a sitting area. This corresponds to the first compartment of the tunnel (40 cm long). The second smaller part of a traditional house is the bedroom where residents sleep under bed netting corresponds to the smaller compartment of the tunnel (20 cm long). We placed the guinea pig within this compartment to represent a sleeping occupant at night (Figure 2). Fifty non-blood-fed female mosquitoes *(An. coluzzii)* were released into the long section of the tunnel. In this design, female mosquitoes are normally attracted through the barrier into the smaller compartment by the guinea pig to blood feed. In full darkness between 6 pm and 6 am, mosquitoes interested in blood feeding were free to fly through the tunnel, locate the holes and pass through them to reach the guinea pig. The location of mosquitoes after this period was recorded, and those in the section closest to the guinea pig were considered to have interest in blood feeding. Mosquitoes were removed from each section of the tunnel and counted separately.

The tunnel bioassays were carried out at 27.34 °C average temperature (range 27.06 °C to 27.60 °C). The average relative humidity was 76.60% (range 75.90% to 77.00%). The mortality during the assay was recorded, but only live mosquitoes were considered for analysis.

#### Bioassay 3: Determination of the effect of *C. violaceum* infection on mosquito fecundity

An overall 75 three days old inseminated females mosquitoes were exposed to *C. violaceum* at 10^6^ bacterial cells / ml according to the infection procedure described above. Control batches of inseminated female mosquitoes were exposed to blank sugar meal without any bacterial cells. For control and treated mosquitoes, as field mosquitoes of this species do not mate in captivity, after adult emergence, inseminations using a force-mating technique between virgin males and females were done according the detail protocol described in MR4 2014 [22]. Control and treated female mosquitoes received then two blood meals, 48hours and 72 hours after insemination. As regard the blood feeding on rabbit, we have ensured that the abdomens of mosquitoes are filled with blood. After the two blood meals, each female was transferred in an individual small cup containing a blotting paper. A layer (~1 cm) of tap water has stayed on the roof of the paper in order to promote eggs laying and hatching. The number of eggs laid per mosquito was scored for the following seven day. The larvae of first instars from eggs were also counted 2-3 days after hatching.

#### Ovary dissection

We therefore attempted to look at the impact of *Chromobacterium violaceum* on the development within both infected inseminated females and uninfected mosquitoes. After the blood meals as above, inseminated females were anesthetized onto cold (−20°C) for five to ten minutes. Then we dissected individually their ovaries under a microscope as described in detailed in MR4 [20]. The aspects of the ovaries were examined under microscope at 40 times magnification. Leica software (LAS-EZ-V3-3-0 for PC) was used to take pictures of the aspects of ovaries

### Data analysis

All data were entered into Microsoft Windows Excel 2013, checked for accuracy, then imported to R studio version 2.11.1 for data manipulation, visualization and statistical analysis (**Supplemental file 2**). Using Fisher’s exact test, P<0.05 was accepted as statistically significant.

The main parameters were calculated at each time point as below:

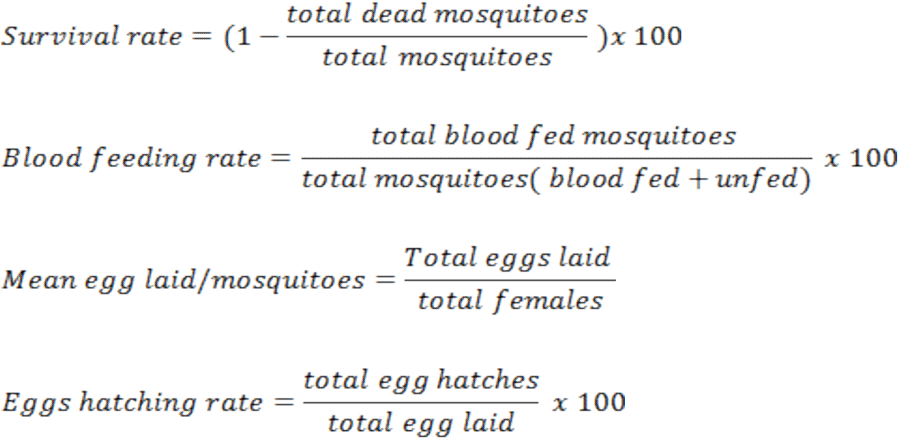

For these main parameters the standards errors (SE) for all replicates were calculated using the library plyr of R 3.2.4:

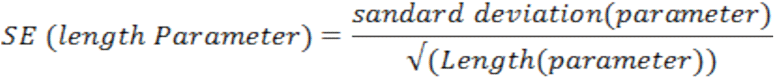

LT_80_ survival for treatments and concentrations were determined using generalized linear model (GLM) approach.

Pairwise t. test comparisons with correction of ohm were used to compare the mean number of mosquitoes per treatments that blood fed, the number of eggs laid and the number of egg hatched respectively. For all bioassays, mosquitoes were considered alive if they could stand upright and dead if they were unresponsive to stimuli following the 2013 recommendations by the WHO Pesticides Evaluation Scheme [23].

#### Ethics statement

All experiments with guinea pigs and rabbits were carried out in strict accordance with the recommendations in the Guide for the Care and Use of Laboratory Animals of the National Institutes of Health [24]. In addition, the protocols followed the IRSS Animal Welfare Assurance A5926-01. Trained personnel and veterinarians cared for animals involved in this study and all efforts were made to minimize suffering. All works with *C. violaceum* were performed under biosafety containment level II requirements.

## Results

### Entomopathogenic effect of C. violaceum on Anopheles coluzzii survival

Overall, 400 female mosquitoes of *An. coluzzii* were infected with serial dilution concentrations of *C. violaceum* from 10^8^ to10^5^ bacterial cells / ml. The level of *kdr* resistance in those wild-caught *An. coluzzii* that we used for bioassays was 98.3%. Within ~9 days post-infection, more than 80% of mosquitoes exposed to the higher concentration to 10^8^ bacterial cells / ml were dead, so significantly faster (P<0.05) than those exposed to the 3 lower concentrations (Figure 3, table 1). However there is no difference (P=0.22) in term of virulence (LT_80_) between 10^7^ and 10^6^ bacterial cells / ml with LT_80_ value of 10.16±0.51 and 10.85±0.42 days respectively (table 1). The lower concentration 10^6^ bacterial cells / ml did not reach the LT_80_ threshold out of 13 days. Observing the survival over 2 weeks, mosquitoes of uninfected control group never dropped below 84.2% (Figure 3).

**Figure 3:**
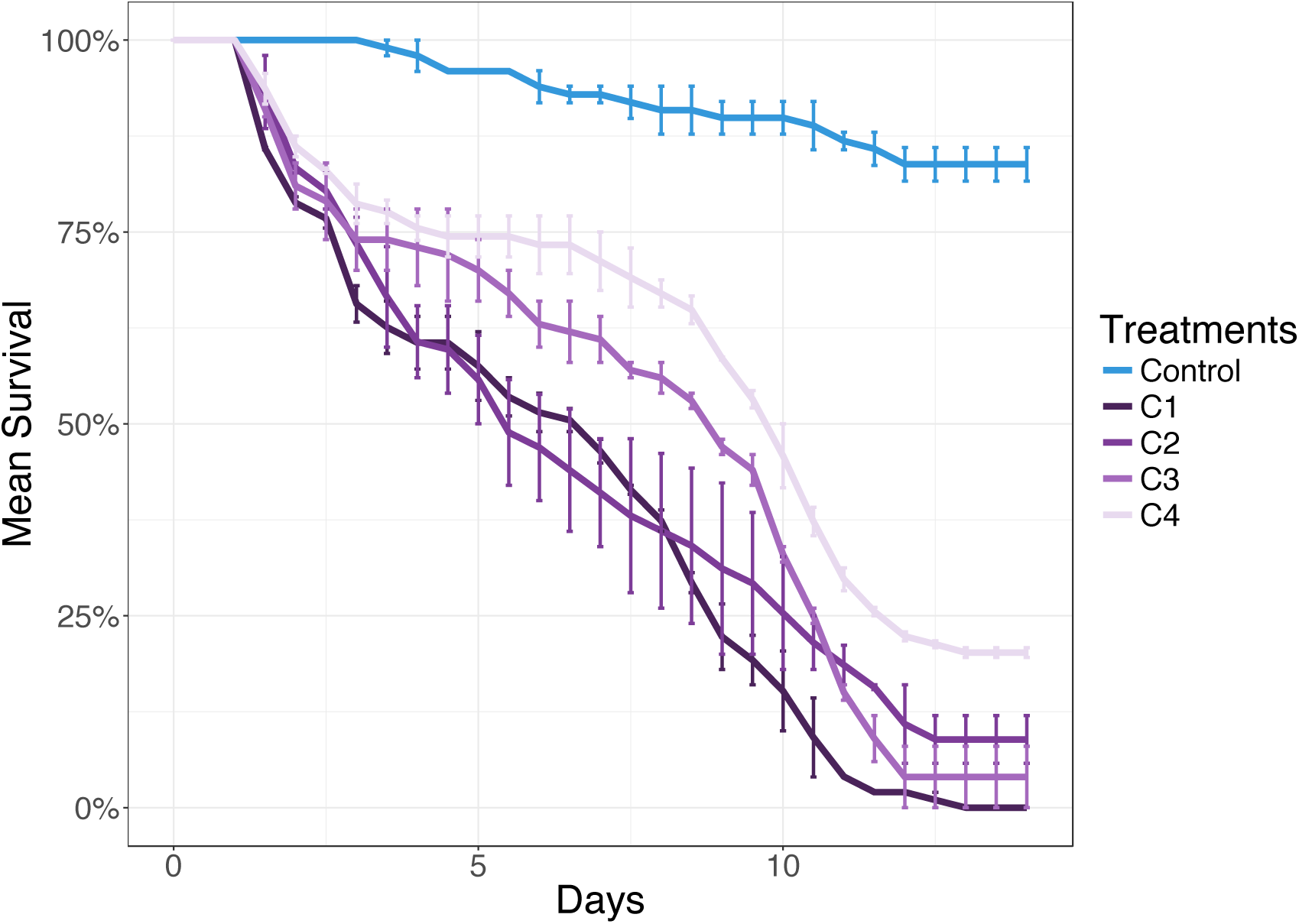
Survival curves of mosquitoes exposed to different concentrations of *C. violaceum*

**Table 1:** LT_80_ survival values of mosquito treated with *C. violaceum*

| Treatments | LT<sub>80</sub> Mean<br>(Days) | S.E.<br>(Days) | Grouping<br>LT<sub>80</sub><sup>2</sup> |
| --- | --- | --- | --- |
| Control | --- | --- | --- |
| C1 | 8.78 | 0.12 | a |
| C2 | 10.16 | 0.51 | b |
| C3 | 10.85 | 0.42 | b |
| C4 | 13.46 | 0.01 | c |

**Legend:** SE: Standard error of the mean; ^2^Pairwise comparison of LT_80_ values per spraying conidia suspension concentrations: Treatments without letters in common are significant at p<0.05. C1, C2, C3 and C4 are 1 × 10^8^ bacteria cells /mL, 1 × 10^7^ bacteria cells /mL, 1 × 10^6^ bacteria cells /mL and in 1 × 10^5^ bacteria cells /mL in 6% glucose meal, respectively. Control is exposed to blank cotton balls soaked 6% glucose meal (without any bacterial cells).

### Impact of C. violaceum infection on females Anopheles coluzzii blood feeding propensy and malaria transmission

Willingness and ability to blood feed were also tested. Host-seeking (blood feeding) interest was quantified as the percentage of the mosquito population choosing to enter and bite the host (guinea pig). At three day post-infection, ~81% of untreated (controls) and treated (*C. violaceum*) mosquitoes flew toward the guinea pig and took their blood meal with no significant differences between treatments. The willingness of mosquitoes in the control group to blood feed did not change over the course of the experiment (from day 3 to day 9 post infection), figure 4.

**Figure 4:**
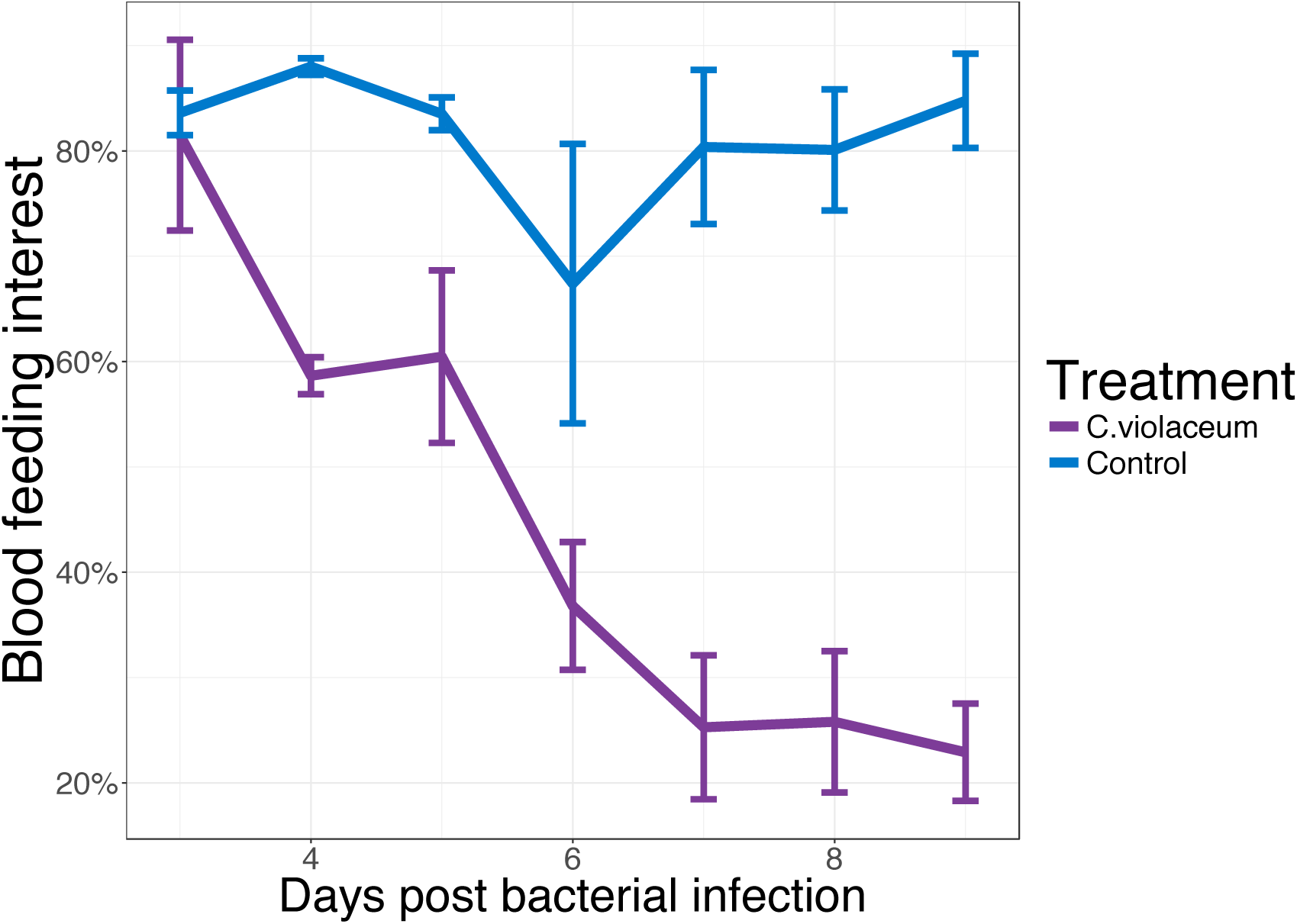
Impact of bacterial infection on blood feeding at 3–9 days post-infection with C. violaceum

In contrast, significantly (p < 0.05) fewer mosquitoes treated with *C.violaceum* flew and had the blood meal after from day 4-post infection to day 9-post infection as compared to Control (P<0.05), figure 4. From day 4 to 9 days post infection we observed an important reduction (59%) of blooding interest with mosquitoes treated with *C. violaceum*. By nine-day post infection the number of *C. violaceum* infected mosquitoes proportion in the guinea pig choice chamber (22±4.62%) was not significantly different than the 30% entering the chamber in the absence of a guinea pig.

The results above suggest a pre-lethal advantage of using *C. violaceum* for mosquito control. With this information, we projected the measured proportion of mosquitoes interested in blood feeding onto the mortality of mosquitoes between 3 and 7 days post infection in order to identify the fraction capable of malaria transmission (Figure 5). In contrast to untreated mosquitoes, by day 6, *C. violaceum* infected mosquitoes passed the threshold in both metrics (18.95 % malaria transmission and >80% mortality) for the 80% control threshold suggested by the World Health Organization (WHO) for a successful vector control agent.

**Figure 5:**
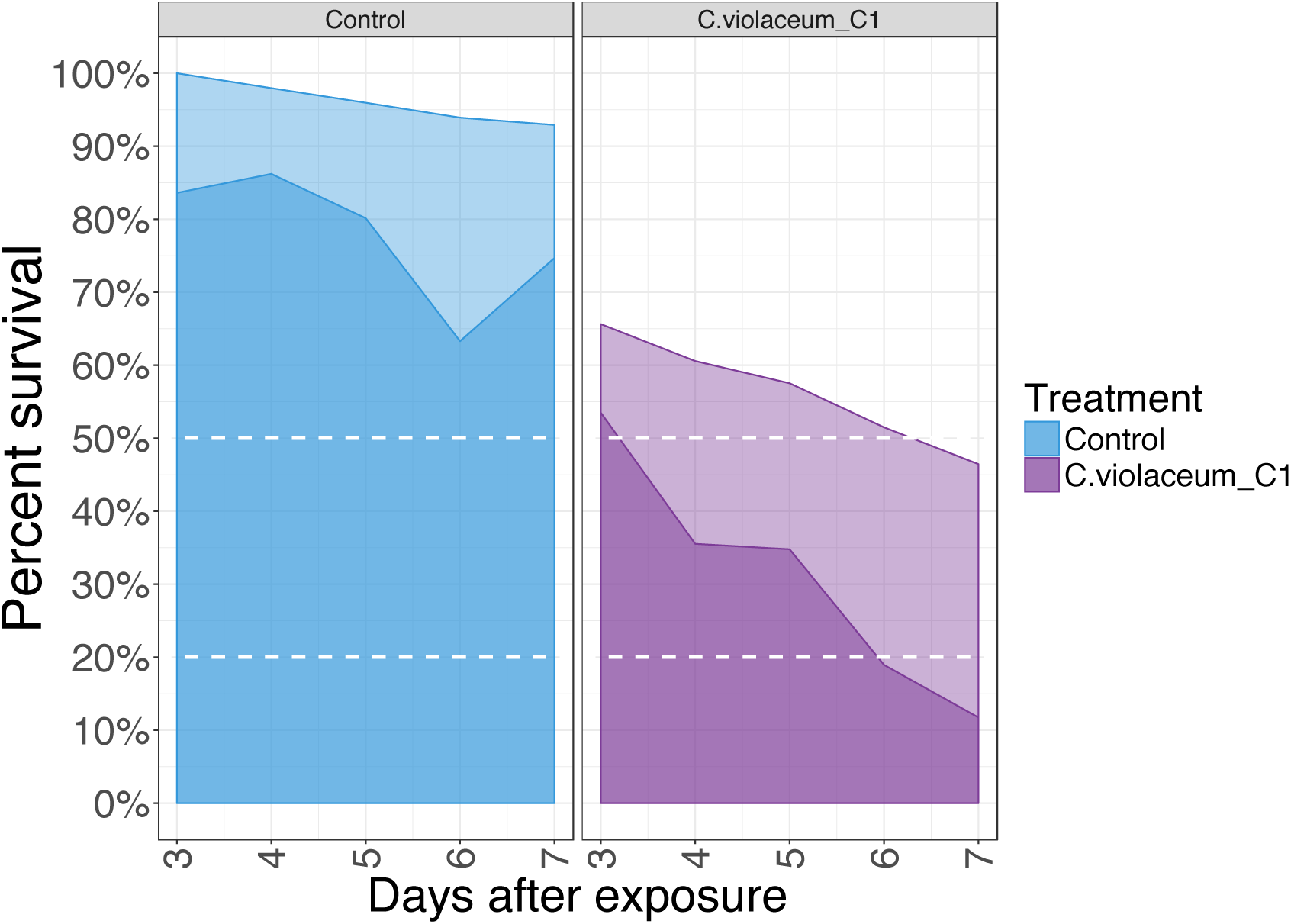
Mortality and transmission of mosquitoes exposed *to C. violaceum*

#### Impact of C. violaceum infection on females Anopheles coluzzii fecundity

The figure 6 shows the pattern of the number of eggs laid by 75 inseminated females infected with *C. violaceum* and 75 inseminated non-infected females (Control).

**Figure 6:**
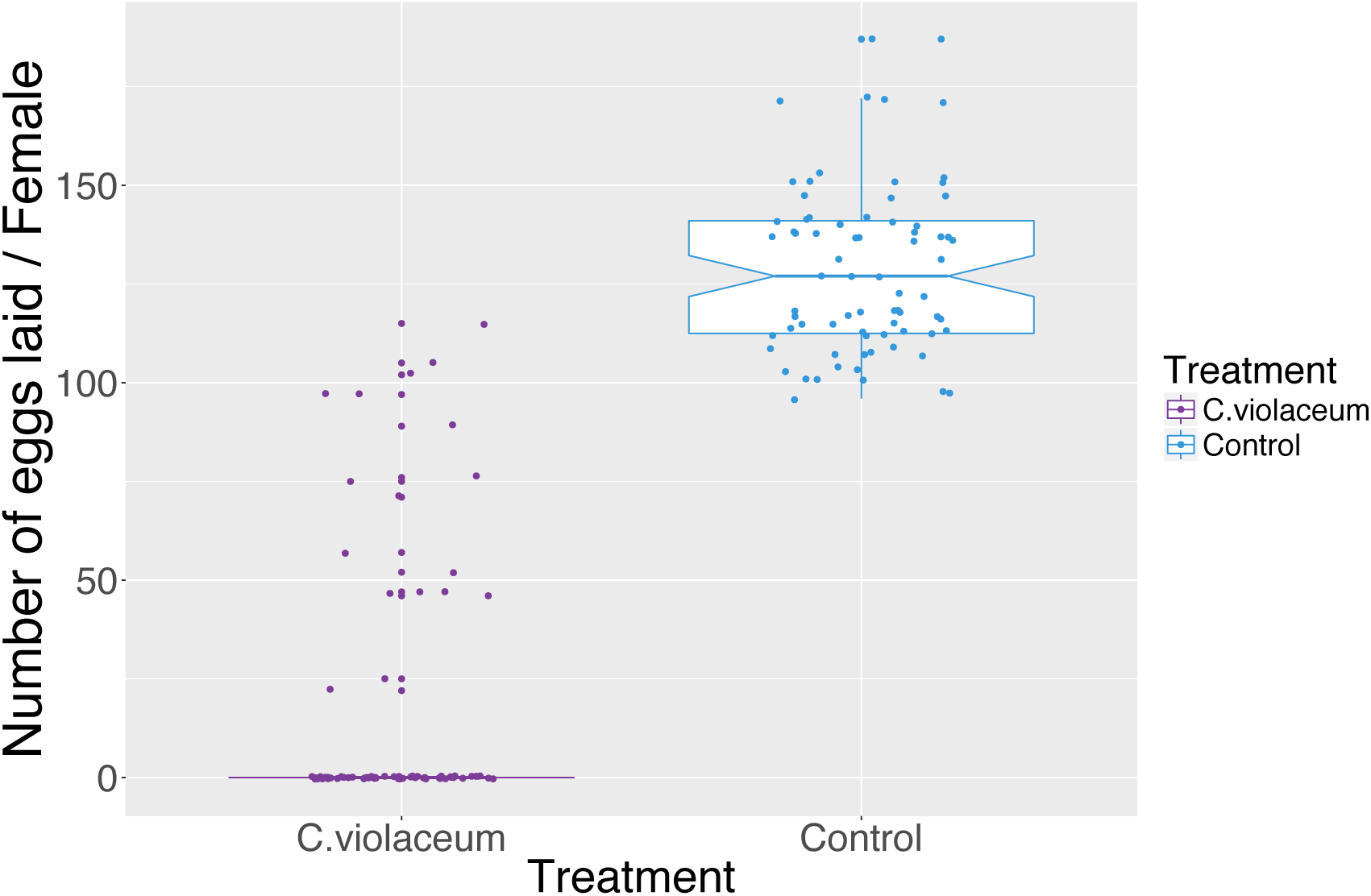
Impact of bacterial infection (*C. violaceum*) on mosquitoes (*An. coluzzii*) egg laying propensity

Overall, 1,170 eggs were laid by 17 females in the *C. violaceum* treated group while 9,648 eggs were laid by 75 females in the control group. Between *C. violaceum* treated mosquitoes and uninfected mosquitoes, we observed a significant difference in terms of mean egg laying propensity per female (P<0.001). A microscopic examination of the aspects of ovaries of inseminated females infected with *C. violaceum* showed a colonization of ovaries of females by *C. violaceum* leading to eggs and ovarian deformities (Figure 7).

**Figure 7:**
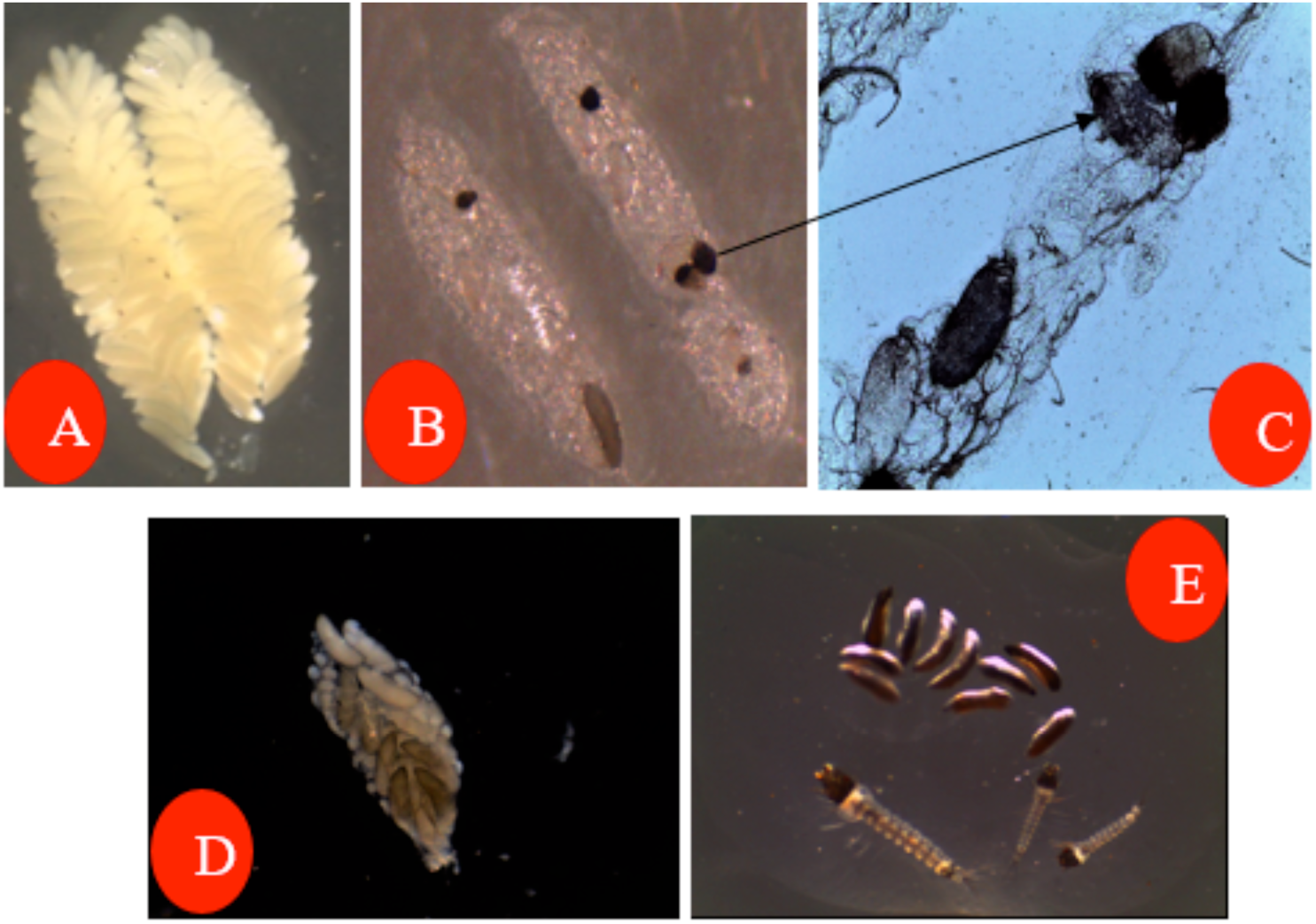
Impact of C. violaceum infections on ovarian follicles and fertilized egg maturations in *An. Coluzzii* mosquitoes.

**Legend**: Eggs of an uninfected female (A); Follicles and fertilized eggs of infected female with C.violaceum (B-D); non-viable eggs and larvae of an infected female (E).

**Figure 8:**
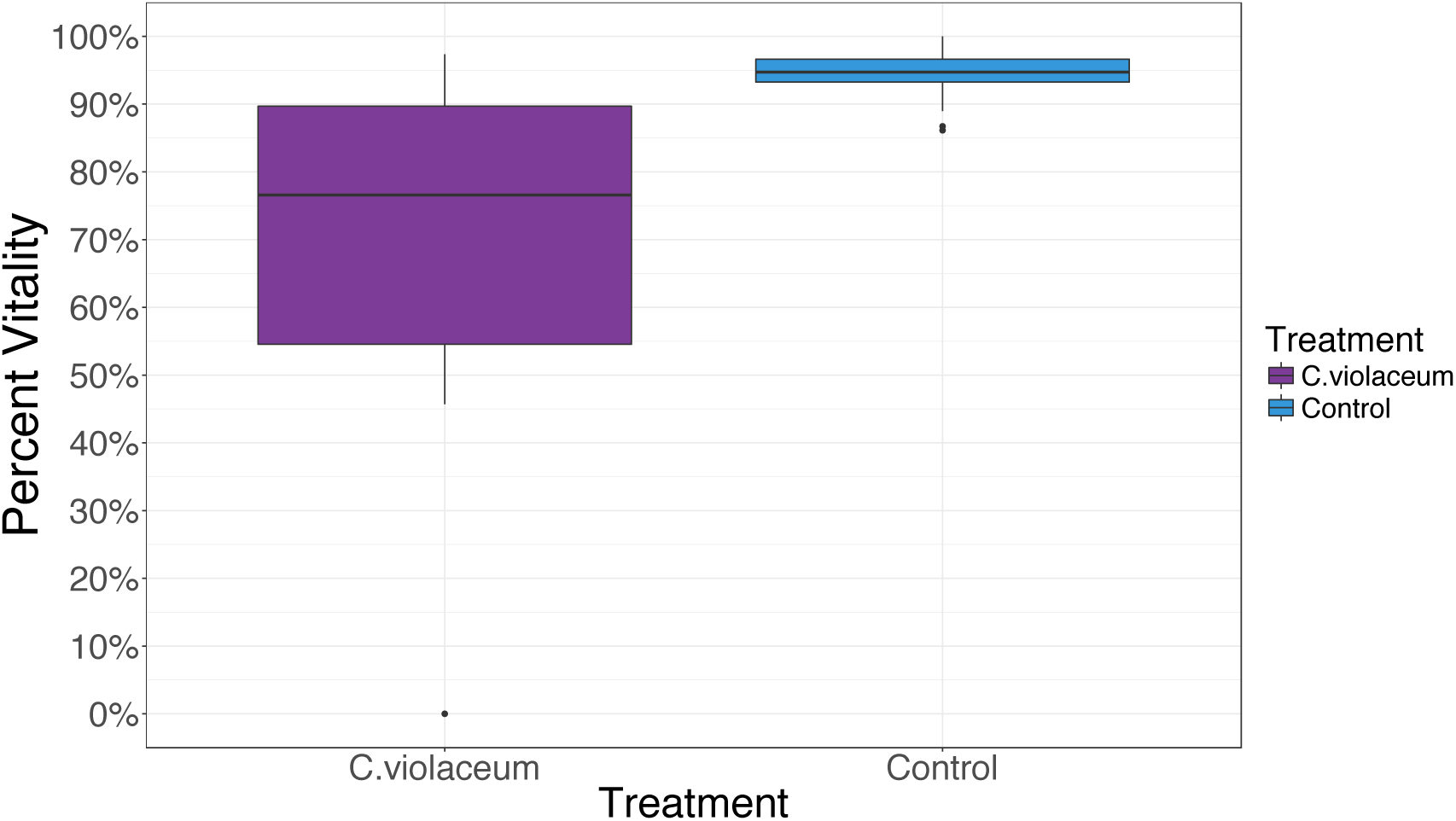
Effect of bacterial infection (*C. violaceum*) on larval vitality rates in mosquitoes (*An. coluzzii*)

Regarding the hatching rate and vitality of first instar larvae from the two groups, we also found significant reduction of ~22% in the *C.violaceum* treated mosquitoes (Figure 8).

## Discussion

*Chromobacterium violaceum* has shown an oral toxicity in highly pyrethroides malaria vector *An. coluzzii*. With a medium concentration, this local strain of *Chromobacterium* surpassed the mortality of 80% thresholds of WHO over a week. *Chromobacterium violaceum* exerted an important entomopathogenic activity even the presence of other microorganisms from mosquito microbiota. This strain of Chromobacterium, unlike the Ramirez et *al.’s* work in 2014 that had isolated *Chromobacterium sp* from *Aedes aegypti* midgut [10]. This strain was isolated from larvae and adults of *An. gambiae s.l*. in western Burkina Faso. Our virulence results corroborate those found by Ramirez et al. in 2014 that also showed a low survival rate of *An. gambiae* and *Ae. aegypti* after a blood meal containing *Chromobacterium sp* at 10^8^ bacterial cells / ml. However the entomopathogenic effect of Chromobacterium species upon *Anopheles gambiae s.l*. is still to be elucidated. A number of potential virulence factors may contribute to mosquitocidal effect, including production of the pigment violacein, siderophores, hydrogen cyanide, and secreted chitinases [25]. In addition, some strains Chromobacterium are capable of forming biofilms in vitro, though whether biofilm formation occurs within the mosquito midgut remains untested [10]. Bacterial biofilms are structured clusters of bacterial cells coated with a polymeric matrix and attached to a surface. The biofilm protects bacteria and allows them to survive in harsh environmental conditions and to resist the immune response of the host. The ability to form a biofilm is now recognized as a characteristic of many entomopathogenic microorganisms [26].

Our results indicated that the rapid death of mosquitoes however is only part of the story. Once infected with *C. violaceum*, mosquitoes are less inclined to blood feed which mainly appear a common effect of fungal infection in mosquitoes and non for bacterial infections [21][27].This blood feeding reduction effect appears stronger as the bacterial infection progresses but can contribute to significant reductions in host feeding as early as day 6, essentially accelerating the transmission blocking. What is striking here is that when the effects of blood feeding are added in, risk of malaria transmission is essentially reduced to zero within 4 day of bacterial exposure and never recovers. *C. violaceum* infection in mosquitoes could share the same physiological impact in term of blood feeding reduction as with fungi. Fungal infection actually increases mosquito metabolic rate and reduces flight propensity and flight stamina. So poor flight performance has been strongly associated with reductions in the mobile energy reserves of the host [28, 29]. Further specific studies are needed to access the impact of *C. violaceum* infection on mosquito flight ability associated with blood feeding propensy to withdraw more conclusions.

One of the results that have never been previously shown to our knowledge is the disruption of mosquito reproduction after infection with *C. violaceum*. Bioassays showed that *C. violaceum* inhibited egg development in the ovaries of *An. coluzzii* mosquitoes. Egg exposures to *C. violaceum* appear to negatively impact ovarian follicles development. Female mosquitoes require blood meals their egg maturations. Newly emerged female mosquitoes come out from the larval stage with theirs follicles at first stage (Stage I). They need blood meals to complete the development of their ovaries to the follicle stage V. In our study, 24 hours of exposure to *C. violaceum* reduced and subsequently stopped the maturation of the follicle because they have not reached the stage V for the most of them. *C. violaceum* could secrete a hormone or substance that directly affects the development of the ovaries and eggs. This substance could indirectly inhibit the action of the juvenile hormones (JHs) in mosquitoes. The juvenile hormones are mainly responsible for the development of ovaries and eggs in mosquitoes [30]. The biochemical and physiological mechanisms governing the innervation of mosquito fecundity following *C. violaceum* infection remain to be elucidated. On other hand, a few infected females of *An. coluzzii* that were able to lay a couple of eggs were inviable or had a low hatching rate. They were infected by *C. violaceum* infection shares this vertical transmission with some bacteria as *Serratia sp, Pantoa agglormerans, Asaia* sp [14, 31–34]. However we need to test if *C. violaceum* shares the cytoplasmic incompatibility properties as some strains of *Wolbachia* [35]. In the simplest case, we can bioassay crosses between *C. violaceum* infected and uninfected *Anopheles coluzzii* mosquitoes in order to see if early embryonic arrest occurs when uninfected females mate with infected males.

## Conclusion

There is a global consensus that in the fight against malaria a ‘magic bullet’ does not exist. The disease can only be controlled by the coordinated deployment of as many weapons as possible. Here, we have isolated and identified a *Chromobacterium violaceum* strain from *Anopheles gambiae* s.l. which exerted an important entomopathogenic effect upon mosquito ingestions. Moreover this strain gives an interesting possibility that it rapidly makes mosquitoes sick, reduces its blood feeding willingness and disrupting its reproduction propensy. These unique properties of *C. violaceum* combined with its environmentally friendly and logistically simple considerations could render this strain a highly potent malaria vector control biopesticide. It should be noted that further investigations are required in other to withdraw more conclusion on the efficacy of *C. violaceum* to control malaria vector. The strain of *Chromobacterium violaceum* although identified to level of species the Vitek 2 compact automated system, it would be important to proceed to the sequencing of the genome of this bacterium and compare it with other species of *Chromobacterium spp*. This will certainly learn us on the specificity of this strain and the role of some of genes in its properties. Regarding the mechanisms that govern properties of *C. violaceum*, bioassays on the impact the bacterial infection on mortality, blood feeding and fecundity propensy on aseptic *Anopheles coluzzii*.

## • Availability of data and material

The datasets generated during the current study is in supplementary file 3. Please, for any request of data or strains contact Abdoulaye Diabate at the Email address: or Etienne Bilgo at the Email address

## List of abbreviations

***C. violaceum***: *Chromobacterium violaceum*
**LT**_50_: is the median (50%) Lethal Time (time until death) after exposure of a mosquito to bacterial infections.
LT_80_: is the 80% Lethal Time (time until death) after exposure of a mosquito bacterial infections

## Declarations

### • Ethics approval and consent to participate

Experiments with animals were carried out in strict accordance with the recommendations in the Guide for the Care and Use of Laboratory Animals of the National Institutes of Health. In addition, Experiments followed the IRSS Animal Welfare Assurance A5926-01. Trained personnel and veterinarians cared for animals involved in this study and all efforts were made to minimize suffering. All works with *C. violaceum* were performed under biosafety containment level II requirements.

### • Consent for publication

All authors have approved the final manuscript and consent for the publication

### • Authors’ contributions

A.D., E.B., A.S., R.K.D. and E.J.G. designed the experiments,

E.J.G. and E.B. performed the experiments and analyzed the data

E.B. and E.J.G. wrote the manuscript.

A.D. is guarantor of the study.

All authors have read and approved the final manuscript.

## • Acknowledgements

We are very grateful to Gnada Kobo Daniel, Eli Kabré, Athur Djibougou, Ouattara Abel Kader, Saré Issiaka for their technical contributions to the field and lab works

## • Endnotes

Not applicable

